# Handedness is associated with less common input to spinal motor neurons innervating different hand muscles

**DOI:** 10.1101/2022.05.25.493122

**Authors:** Jean Maillet, Simon Avrillon, Antoine Nordez, Jeremy Rossi, François Hug

**Affiliations:** Nantes Université, Movement - Interactions - Performance, MIP, UR 4334, F-44000 Nantes, France; Neuromechanics & Rehabilitation Technology Group, Department of Bioengineering, Faculty of Engineering, Imperial College London, UK; Institut Universitaire de France (IUF), Paris, France; Univ Lyon, UJM-Saint-Etienne, Laboratoire Interuniversitaire de Biologie de la Motricité, EA 7424, F-42023 Saint-Etienne, France; Université Côte d’Azur, LAMHESS, Nice, France; The University of Queensland, School of Biomedical Sciences, Brisbane, QLD, Australia

**Keywords:** dominance, common drive, electromyography, motor unit, coherence

## Abstract

Whether the neural control of manual behaviours differs between the dominant and non-dominant hand is poorly understood. This study aimed to determine whether the level of common synaptic input to motor neurons innervating the same or different muscles differs between the dominant and the non-dominant hand. Seventeen participants performed two motor tasks with distinct mechanical requirements: an isometric pinch and an isometric rotation of a pinched dial. Each task was performed at 30% of maximum effort and was repeated with the dominant and non-dominant hand. Motor units were identified from two intrinsic (flexor digitorum interosseous and thenar) and one extrinsic muscle (flexor digitorum superficialis) from high-density surface electromyography recordings. Two complementary approaches were used to estimate common synaptic inputs. First, we calculated the coherence between groups of motor neurons from the same and from different muscles. Then, we estimated the common input for all pairs of motor neurons by correlating the low-frequency oscillations of their discharge rate. Both analyses led to the same conclusion, indicating less common synaptic input between motor neurons innervating different muscles in the dominant hand than in the non-dominant hand, which was only observed during the isometric rotation task. No differences in common input were observed between motor neurons of the same muscle. This lower level of common input could confer higher flexibility in the recruitment of motor units, and therefore, in mechanical outputs. Whether this difference between the dominant and non-dominant arm is the cause or the consequence of handedness remains to be determined.

**Key points:**

- How the neural control of manual behaviours differs between the dominant and non-dominant hand remains poorly understood.
- We decoded the spiking activities of spinal motor neurons innervating one extrinsic and two intrinsic hand muscles during isometric tasks.
- We estimated the common synaptic input to motor neurons innervating the same or different muscles.
- There is less common synaptic input between motor neurons innervating different muscles in the dominant than in the non-dominant hand during isometric rotation tasks.
- No differences in common input were observed between motor neurons of the same muscle.
- Lower level of common input could confer higher flexibility in the recruitment of motor units.

## 1. Introduction

Handedness (or laterality) is a human attribute defined by a preference for the use of one hand, over the other, which is defined as the preferred or dominant hand. Humans almost always use their preferred hand to manipulate an object (for example, a hammer) and their non-preferred hand to stabilise an object (for example, a nail) (Wiesendanger *et al*., 1996). However, how the neural control of manual behaviours differs between the dominant and non-dominant hand remains poorly understood.

The human hand can perform a large repertoire of movements that are fundamental for our interaction with the environment. The versatility of function of the hand is made possible by a complex neuromusculoskeletal system, which involves 31 muscles (including 19 intrinsic muscles), 18 tendons crossing the wrist, and at least 25 degrees of freedom (van Duinen & Gandevia, 2011). This system renders the control of even simple hand movements, such as grasping an object or opening a bottle, complex at both the physiological and the computational level. It has been proposed that the extrinsic and intrinsic hand muscles acting on the fingers are controlled through a modular organization of descending neural inputs, referred to as ‘synergies’ (Weiss & Flanders, 2004; Ajiboye & Weir, 2009; Santello *et al*., 2013). This modular organization implies that groups of spinal motor neurons receive common synaptic inputs (Tanzarella *et al*., 2021), which ultimately reduces the dimensionality of movement control.

Previous studies have assessed the effect of handedness on the common synaptic inputs to spinal motor neurons that innervate intrinsic and extrinsic hand muscles. Specifically, they have calculated the correlation of motor unit discharge times (short-term synchronization) for pairs of motor units from the same pool (Kamen *et al*., 1992; Schmied *et al*., 1994; Semmler & Nordstrom, 1995) or from different muscles (Hockensmith *et al*., 2005). These studies have yielded mixed results, with either greater (Kamen *et al*., 1992; Schmied *et al*., 1994) or lower (Semmler & Nordstrom, 1995) synchrony between motor units from the same pool on the dominant hand. In addition, Hockensmith et al. (2005) found that the synchrony between motor units from two extrinsic hand muscles was markedly higher in the dominant than in the non-dominant hand. In addition to limitations of the synchronization approach to quantify the strength of common synaptic input between motor neurons (Farina & Negro, 2015), the divergent results can originate from the different muscles being investigated, that is, extrinsic (Schmied *et al*., 1994) or intrinsic hand muscles (Semmler & Nordstrom, 1995). It is also important to note that these studies involved very low-intensity isometric contractions during which participants had to maintain the firing of a couple of motor units (Schmied *et al*., 1994; Semmler & Nordstrom, 1995; Hockensmith *et al*., 2005), which imposed specific constraints, different from those observed during purely force-matched tasks at a higher contraction intensity.

The aim of the present study was to determine whether the level of common synaptic input to motor neurons innervating the same or different muscles differs between the dominant and the non-dominant hand. We first estimated this common input at the muscle level (within and between muscle) by calculating the coherence between groups of motor neurons (Del Vecchio *et al*., 2019; Avrillon *et al*., 2021). Then, we estimated the common input for all pairs of motor neurons through the correlation between the low-frequency oscillations of their discharge rate (De Luca & Erim, 1994; Semmler *et al*., 1997). Motor units were identified from two intrinsic hand muscles (the flexor digitorum interosseous [FDI] and the abductor pollicis brevis referred to as thenar muscle) and one extrinsic hand muscle (the flexor digitorum superficialis [FDS]) during two different motor tasks with distinct mechanical requirements, specifically, isometric pinch and isometric rotation of a pinched dial. While the isometric pinch task mainly required a flexion of both the thumb and the index finger in one main axis, the rotation task involved an abduction and flexion of the thumb, together with an abduction of the index finger. This relatively higher complexity of the rotation task may impose greater neural constraints. Based on a previous study reporting a lower coherence in pianists compared to control participants (Semmler *et al*., 2004), we hypothesised that the dominant hand would exhibit lower common synaptic input within and between muscles. This lower level of common input might confer higher flexibility in the recruitment of motor units, and therefore, in mechanical outputs. Furthermore, because of its higher complexity, we hypothesised that the difference between the arms in terms of common input would be mainly observed for the rotation task.

## 2. Methods

### 2.1. Participants

Seventeen healthy volunteers participated in this study (14 males and three females; age: 25.3±4.1 years; height: 1.78±0.09 m; body mass: 69.4±9.2 kg). Fourteen participants were right-handed (laterality quotient: 86.3±11.7) and three were left-handed (laterality quotient: - 54.9±31.7), as tested with the Edinburgh Handedness Inventory (Oldfield, 1971). They had no history of upper limb pain resulting in limited function that required time off work or physical activity, or a consultation with a health practitioner in the previous 6 months. The local ethics committee approved the study (CERNI approval number: 20032020-1), and all participants provided informed written consent.

### 2.2. Experimental design

Each participant enrolled in the study participated in an experimental session lasting ∼3 hours. After a series of motor function tests, the participants were asked to perform two motor tasks (an isometric pinch and an isometric rotation of a pinched dial). These motor tasks were repeated with the right and the left hand, in a randomised order.

#### 2.2.1. Motor function tests

To assess the difference in motor function between hands, participants performed a series of motor function tests. First, they performed a dexterity test, the Purdue Pegboard Test (Tiffin & Asher, 1948). The test required participants to place as many metal pegs as possible into holes on a board within 30 s. The testing board consisted of a board with four cups across the top and two vertical rows of 25 small holes down the centre. Performance was quantified as the total number of pegs placed in the right/left column using the right/left hand in the allocated time. The test was repeated three times for each hand (randomised order) with 30-s of rest in between. The best performance was retained for further analysis.

Second, participants performed two-finger tapping tests to assess their motor speed (Herve *et al*., 2005): one with the index finger and another one with the index and middle finger (alternate tapping). For each task, the participants were instructed to tap as quickly as possible on a mobile phone within a 10-s time interval. The performance was recorded with an application for mobile phone (Click Speed Test, Mnt Apps, USA). The participant’s palm and the other fingers were to rest on the table. Each test was repeated three times for each hand (randomised order) with 30-s of rest in between. The best performance was retained for further analysis.

#### 2.2.2. Experimental tasks

The experimental tasks consisted of either pinching a force sensor isometrically (referred to as the *pinch* task) or applying an isometric rotation force on a custom-made octagonal dial attached to a force sensor (referred to as the *rotation* task). The rotation task was performed anticlockwise with the right hand and clockwise for the left hand, such that the motor task was the same between sides. The index finger and the thumb held either the force sensor (thickness: 4 cm) or the octagonal dial (diameter: 2.5 cm) with the pulp of their distal phalanx. All fingers joints were slightly flexed. For both tasks, the participants were seated with their elbow flexed at 45° (0° = full extension), and their forearm was supported by a comfortable pad to isolate the function of the hand. The wrist and the hand were in a neutral position. Participants were given time to familiarise themselves with each experimental task before starting the protocol described below.

The participants performed both tasks with each hand (randomised order). For each task, they started with a warm-up composed of 10 submaximal contractions. Then, they performed three maximal isometric contractions for 3-s with a 60-s rest in between. The maximal value obtained from a moving average window of 250-ms was considered as the peak force (MVC). Then, the participants performed three contractions involving a 5-s ramp-up, a 20-s plateau at 30% of MVC, and a 5-s ramp-down phase. The contractions were separated by 60-s of rest. This protocol was repeated twice, once on each side. Feedback from the target and force output was displayed on a monitor. Participants were monitored to ensure that they only used their index and their thumb to perform the task. Force and torque signals from the six-axes sensor (Nano25-E, ATI Industrial Automation, USA) were digitised at 2048 Hz. The variability of force (pinch) or torque (rotation) was calculated as the mean coefficient of variation between the three plateaus.

### 2.3. High-density surface electromyography recordings

High-density EMG (HDsEMG) signals were recorded from one extrinsic muscle (FDS) and two intrinsic muscles (FDI, and abductor pollicis brevis). Of note, despite we aimed to measure the abductor pollicis brevis muscle, we cannot ascertain that we selectively measured this muscle. Therefore, we considered that we measured muscles from the thenar eminence, referred to as thenar muscles thereafter. The FDI is an abductor and flexor of the index and an adductor of the thumb. The thenar muscles are abductors and flexors of the thumb. The FDS is a flexor of the long fingers that includes the index finger.

Two-dimensional adhesive grids of 64 electrodes (13×5 electrodes with one electrode absent on a corner, gold-coated, inter-electrode distance: 4 mm; [GR04MM1305; OT Bioelettronica, Italy]) were placed on the thenar muscle and FDI as described in Del Vecchio et al. (2019). Another grid (13×5 electrodes with one electrode absent on a corner, gold-coated, inter-electrode distance: 8 mm; [ELSCH064NM2; SpesMedica, Italy or GR08MM1305 OT Bioelettronica, Italy]) was placed over the FDS muscle. To ensure the proper location of this grid, the borders of the FDS muscle were identified using B-mode ultrasound (Aixplorer, Supersonic Imagine, France). Before placing the grids, the skin was shaved, and then cleaned with an abrasive pad and alcohol. The adhesive grids were held on the skin using semi-disposable bi-adhesive foam layers. The skin-electrode contact was made by filling the cavities of the adhesive layers with conductive paste (SpesMedica, Battipaglia, Italy). A reference electrode (Kendall Medi-Trace(tm), Canada) was positioned over the olecranon process of the elbow. A strap electrode dampened with water was placed around the contralateral wrist (ground electrode). The EMG signals were recorded in monopolar mode, bandpass filtered (10– 500 Hz) and digitized at a sampling rate of 2048 Hz using a multichannel HDsEMG acquisition system (EMG-Quattrocento, 400 channel EMG amplifier; OT Bioelettronica, Italy). Once data collection was completed for one hand, the electrodes were moved to the contralateral side.

### 2.4. Data analysis

Data were analysed using MATLAB custom-written scripts (R2021a, The Mathworks, Natick, MA, USA).

#### 2.4.1. HDsEMG decomposition

First, the monopolar EMG signals were bandpass filtered between 20 and 500 Hz with a second-order Butterworth filter. After visual inspection, channels with a low signal-to-noise ratio or artifacts were discarded. The HDsEMG signals were then decomposed into motor unit spike trains. Specifically, we applied a convolutive blind-source separation as previously described (Negro *et al*., 2016). In summary, the EMG signals were first extended and whitened. Thereafter, a fixed-point algorithm was applied that maximised the sparsity to identify the sources of the EMG signals, that is, the motor unit spike trains. The spikes were separated from the noise using K-mean classification and a second algorithm refined the estimation of the discharge times by minimizing the coefficient of variation of the inter-spike intervals. This decomposition procedure has been previously validated using experimental and simulated signals (Negro *et al*., 2016). After the automatic identification of the motor units, all the motor unit spike trains were visually checked for false positives and false negatives (Del Vecchio *et al*., 2020; Hug *et al*., 2021a). All motor units that exhibited a pulse-to-noise ratio > 30 dB were retained for further analysis. An operator (JM) analysed all the data, and the reliability of each motor unit spike train was systematically checked an experienced operator (FH). It should be noted that the manual edition of the motor unit spike trains has been shown to be highly reliable across operators (Hug *et al*., 2021a).

#### 2.4.2. Within- and between-muscle coherence

To estimate the level of common input at the population level, we calculated the coherence between cumulative spike trains (CST) of motor units from the same (within-muscle coherence) or from different muscles (between-muscle coherence) (Fig. 1). The coherence represents the correlation between two signals at given frequencies, with 0 indicating no correlation and 1 indicating a perfect correlation. Coherence within the delta band (0-5 Hz) reflects the presence of common drive (De Luca *et al*., 1982; Del Vecchio *et al*., 2019), while coherence within the alpha band (5-15 Hz) reflects the contribution of the muscle afferents and other spinal circuitries (Williams & Baker, 2009). Of note, we did not focus on the beta band (15-35 Hz), because a larger number of motor units is needed to accurately estimate coherence within this bandwidth. This is mainly due to the non-linear relationship between the synaptic input to motor neurons and their output signal, which is more pronounced at higher frequencies due to an undersampling of the synaptic input (Negro & Farina, 2012; Farina & Negro, 2015).

**Fig 1.**
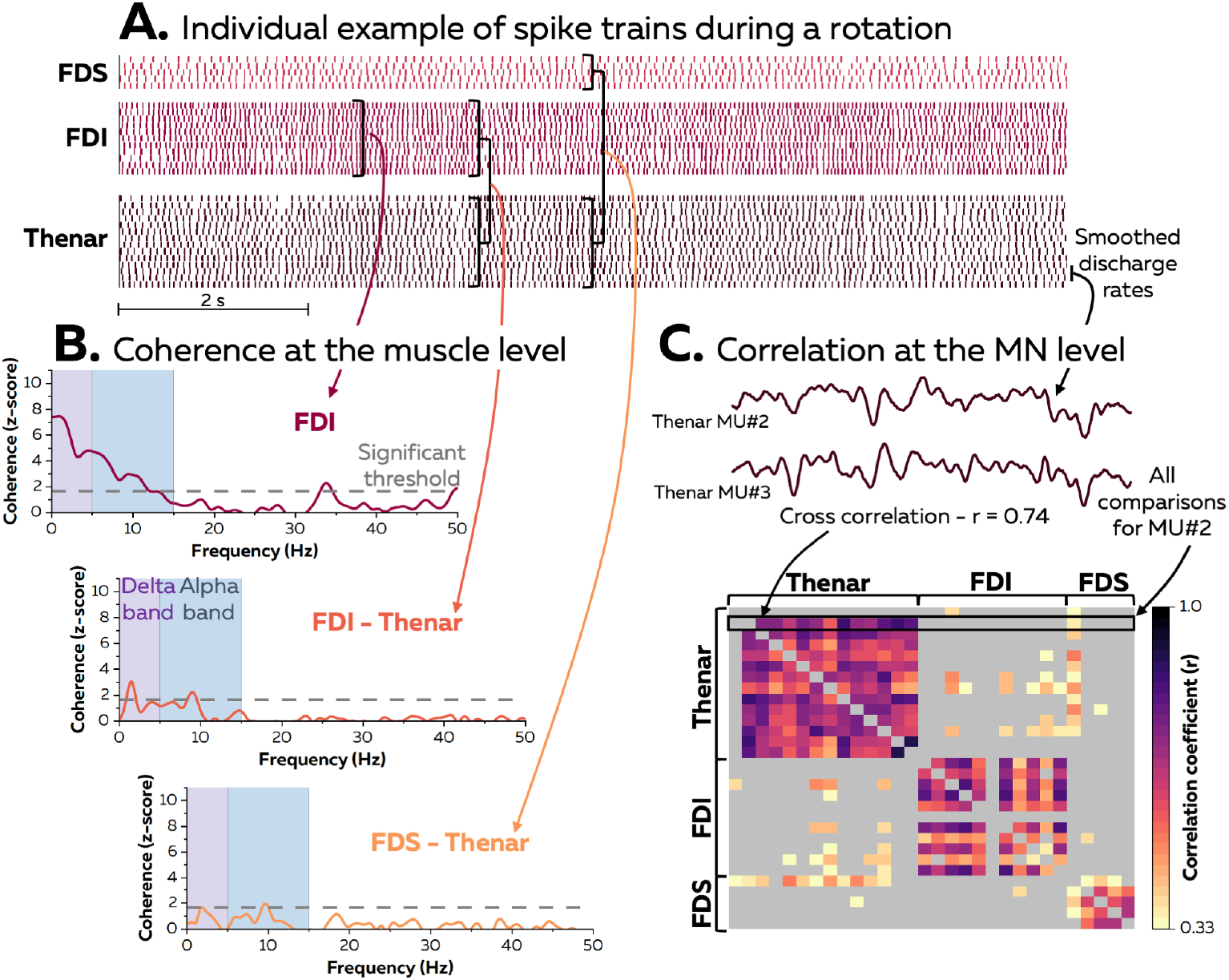
Experimental measures. The decomposed motor unit discharge times were first converted into continuous binary signals (Panel A). We assessed the common input using two complementary approaches. First, we estimated the common input at the muscle level by calculating the within- and between-muscle coherence (Panel B). Second, we estimated the common input for each pair of motor neurons. To this end, a cross-correlation function was applied to the smoothed discharge rates, and only the correlation coefficients that reached a significant threshold of 0.33 (see Methods) were considered (Panel C). FDI, first dorsal interosseous; FDS, flexor digitorum superficialis; MN, motor neuron.

Only the plateau of the trapezoid contractions was considered for the coherence analyses. To assess the coherence within and between muscles, the magnitude-squared coherence was calculated using the Welch’s averaged periodogram with nonoverlapping windows of 1-s over a 30-s time window, which was a concatenation of multiple windows from the three contractions. These windows were selected such that the number of active motor units was maximised. This analysis was performed on two equally sized cumulative spike trains calculated as the sum of discharge times from two motor units randomly selected from the identified motor units from the same muscle (within-muscle coherence) or two different muscles (between-muscle coherence). This was repeated for all the unique combinations of two motor units per group, up to a maximum of 100 random permutations. We retained the pooled coherence of these random permutations for further analysis.

As the level of coherence depends on the number of motor units considered in the analysis (Farina and Negro 2015), we considered a fixed number of motor units (n=2) per group to facilitate comparison between muscles (or muscle pairs), tasks, and sides. The number of motor units per group was chosen with the intention to maximise the number of participants for whom these analyses were possible. Despite this, only a few participants exhibited constant firings of at least four units (i.e. two groups of two units) within a 30-s time window for the thenar and FDS muscle. Therefore, within-muscle coherence for these two muscles is not reported. Similarly, only a few participants exhibited constant firings of at least two units per muscle within a 30-s time window for the FDI-FDS pair. Therefore, between-muscle coherence values for this muscle pair were not included in the statistical analysis.

We transformed both the within and between-muscle coherence values into standard z-scores as described in previous studies (Del Vecchio *et al*., 2019; Avrillon *et al*., 2021):

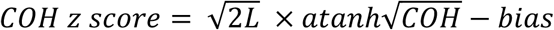

where COH is coherence, L is the number of time segments used for the coherence analysis (e.g. for 30s, L = 30 as the analysis was performed on 30 windows of 1-s), and bias is the mean COH z-score between 250 and 500 Hz where no coherence is expected (Baker *et al*., 2003). Coherence was considered significant when the z-score was greater than 1.65 (95% confidence limit).

#### 2.4.3. Correlation between smoothed motor unit spike trains

To estimate the level of common input between individual motor neurons, we calculated the correlation between their smoothed discharge rates (Fig. 1). To this end, the decomposed motor unit spike times were first converted into continuous binary signals with ‘ones’ corresponding to the firing instances of a unit. To limit the effect of the non-linear relationship between the synaptic input and the output signal, these binary signals were convoluted with a 400-ms Hanning window, such that the correlation was calculated from the low-frequency oscillations of the signal (Semmler *et al*., 1997; Negro & Farina, 2012). Finally, these signals were high-pass filtered with a cut-off frequency of 0.75 Hz to remove offsets and trends. The level of common input for each pair of motor neurons was estimated using a cross-correlation function, and the maximum cross-correlation coefficients with a maximal lag of ± 100 ms were retained for further analysis (De Luca & Erim, 2002). This was performed in a 10-s window that was chosen such that the number of concurrent active motor neurons across the three muscles was maximized.

To account for the fact that the strength of common synaptic inputs between two motor neurons is not proportional to the degree of correlation between their outputs (de la Rocha *et al*., 2007), we only interpreted the correlation coefficients based on their significance (Hug *et al*., 2021b). Specifically, we defined a significance threshold as the 95^th^ percentile of the cross-correlation coefficient distribution generated with resampled versions of the motor unit spike trains. We generated a surrogate spike train for each motor unit by bootstrapping the interspike intervals (random sampling with replacement). This random spike train had the same number of spikes, and the same discharge rate (mean and standard deviation) as the original motor unit spike train. Two iterations of this random procedure were performed, such that each motor unit was associated with two surrogate spike trains and each motor unit pair with four combinations of surrogate spike trains, thereby yielding to 9,402 correlation coefficients for the whole population. This analysis was performed in the same 10-s window that had been used for the main analysis and yielded to a significant threshold of 0.33.

### 2.5. Statistical analysis

All statistical analyses were implemented in RStudio (USA). We first plotted quantile-quantile plots and histograms to assess the distribution of each dataset. Data that deviated from a normal distribution were transformed with functions adapted to the skew of the distribution.

We compared MVC values and the performance of the Purdue Pegboard Test and Finger Tapping Tests between the dominant and non-dominant sides using paired t-tests.

We then used linear mixed effect models (LMM) implemented in the R package lmerTest with the Kenward-Roger method to estimate the denominator degrees of freedom and the p values. We first compared the coefficient of variation of the force/torque calculated over the 20-s force plateaus (fixed effects: task [pinch, rotation] and dominance [dominant, non-dominant], random effect: participants). Then, to assess the effect of handedness on the level of common synaptic input at the muscle level, we compared the mean within-muscle coherence values (delta and alpha band) for the FDI muscle (fixed effects: task [pinch, rotation] and dominance [dominant, non-dominant], random effect: participants). We then compared the mean values of between-muscle coherence (delta and alpha band) for the muscle pairs (fixed effects: muscle pairs [FDI-Thenar, FDS-Thenar], task [pinch, rotation], and dominance [dominant, non-dominant]; random effect: participants).

To assess the effects of handedness on the level of common synaptic input at the motor neuron level, we first compared the ratio of significant correlations between motor neurons of the same muscle (fixed effects: muscle [Thenar, FDI, FDS], task [pinch, rotation], and dominance [dominant, non-dominant]; random effect: participants). Then, we compared the ratio of significant correlations between motor neurons from two different muscles (fixed effects: muscle [FDI-Thenar, FDS-Thenar, FDI-FDS], task [pinch, rotation], and dominance [dominant, non-dominant]; random effect: participants).

When necessary, we performed multiple comparisons using the R package *emmeans*, which adjusts the *p*-values using the Tukey method. Statistical significance was set at 5% (*p* < 0.05). Values are reported as means ± standard deviation (SD).

## 3. Results

### 3.1. Motor function and force/torque variability

The dexterity assessed using the Purdue Pegboard Test was higher for the dominant (17.8±1.6 pegs in 30s) than for the non-dominant hand (16.8±1.3 pegs in 30s, p=0.009). Similarly, the motor speed evaluated using the Finger Tapping Tests was higher for the dominant than for the non-dominant hand (index finder test: 66.1±7.9 vs. 61.1±7.0 taps; p=0.017; index and second finger test: 85.6±26.6 vs. 72.4±20.3 taps; p=0.009).

Although the MVC was not different between the dominant and the non-dominant hand for the pinch task (p=0.19), the MVC of the non-dominant hand was significantly higher than that of the dominant hand for the rotation task (+12.3±19.1%; p=0.013). When considering the coefficient of variation of the force/torque during the plateau of the submaximal contractions, there was a main effect of task (p=0.032), but no effect of dominance (p=0.368), and neither an interaction between task and dominance (p=0.477). Specifically, the coefficient of variation was higher for the rotation (3.4±4.3%) than for the pinch task (1.8±0.5%).

### 3.2. Motor unit analysis

In total, 1263 motor units (FDI: 693, thenar: 322, FDS: 248) were identified, with an average of 10.2±4.1 (FDI), 5.9±2.6 (thenar), and 5.0±2.1 (FDS) motor units per contraction, side and participant. The entire data set (raw and processed data) is available at https://figshare.com/s/ba9a27d0bec76f2f9bbf.

The mean motor unit discharge rate was calculated for each muscle and each task over the force/torque plateau of the trapezoid contractions. A significant main effect of muscle was observed (F(2, 146)=60.379; p< 0.001) (Fig.2). Specifically, the discharge rate was lower for FDS than for the thenar (p<0.001) or the FDI muscle (p<0.001). There was no difference between the thenar muscle and the FDI muscle (p=0.52). There was no other main effect or interaction (all p values > 0.106), indicating that the discharge rate did not differ between sides. As coherence analysis required the selection of motor units that discharged consistently over the same 30-s window, not all the identified motor units could be included in these analyses. Indeed, some motor units were recruited intermittently, and other motor units (mainly from different muscles) did not necessarily discharge during the same time periods. As a result, within-muscle coherence could be assessed for FDI in 14 participants and between-muscle coherence could be assessed in10 participants for both the FDI-thenar and the FDS-thenar pair. The average number of motor units considered for each analysis is shown in Table 1.

**Fig. 2.**
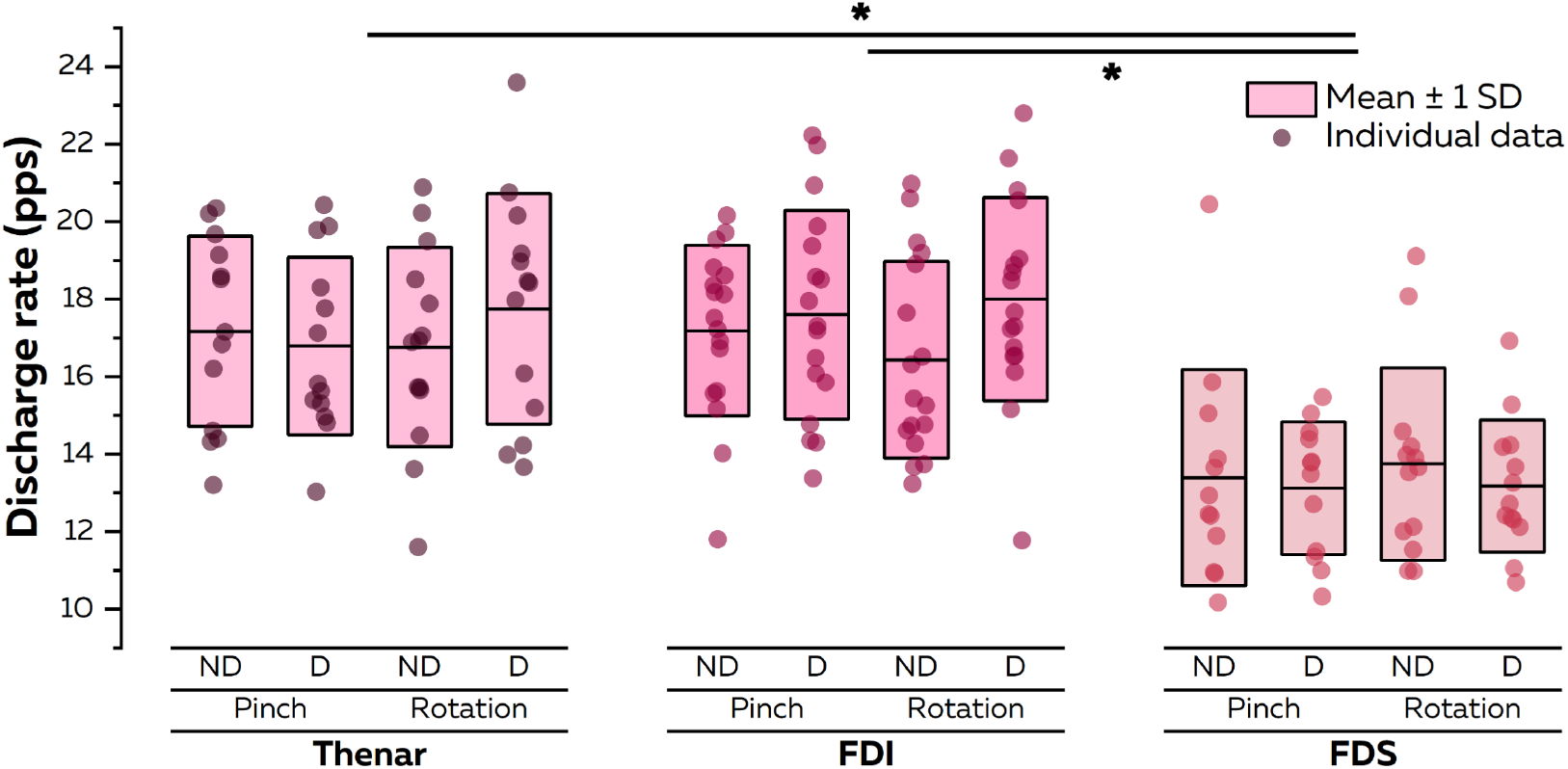
Motor unit discharge rate. The mean motor unit discharge rate was calculated for each muscle and each task over the torque plateau of the three trapezoid contractions. Each participant is represented by a dot. There was a main effect of muscle, with a significant difference between the thenar and the FDS and between the FDI and the FDS. No other main effect or interaction was found. PPS, pulse per second; ND, non-dominant; D, dominant; first dorsal interosseous; FDS, flexor digitorum superficialis.

**Table 1.** Number motor units per participant used for the main analyses. The number of motor units per muscle included in analysis is reported for each muscle/muscle pair (mean ± standard deviation). The number of motor units per muscle is provided in the same order as that for the muscle pair, i.e., first muscle in the left column and second muscle in the right column. FDI, First Dorsal Interosseous; FDS, Flexor Digitorum Superficialis.

|  | Pinch |  | Rotation |  |
| --- | --- | --- | --- | --- |
|  | Non-dominant | Dominant | Non-dominant | Dominant |
| **Coherence analysis** |  |  |  |  |
| FDI | 8.1±2.8 | 7.8±2.9 | 9.1±4.2 | 8.7±3.4 |
| FDI-thenar | 8.6±3.6; 4.2±2.2 | 8.3±3.5; 5.3±1.7 | 9.9±6.5; 4.6±2.9 | 9.9±2.8; 4.8±2.6 |
| FDS-thenar | 3.1±1.8; 3.8±1.5 | 3.1±1.1; 4.4±1.9 | 3.2±1.3; 4.3±2.9 | 3.8±1.3; 4.6±3.0 |
| **Pairwise cross-correlation of smoothed discharge rate** |  |  |  |  |
| FDI | 9.6±3.0 | 8.9±3.8 | 9.8±5.7 | 10.3±3.5 |
| Thenar | 4.5±2.0 | 5.7±1.8 | 5.9±2.5 | 5.3±3.2 |
| FDS | 3.5±1.9 | 3.8±1.8 | 4.3±1.4 | 3.9±1.4 |
| FDI-thenar | 9.6±3.0 – 4.5±2.0 | 8.9±3.8 – 5.3±2.1 | 9.8±5.7 – 5.5±2.8 | 10.3±3.5 – 5.3±3.2 |
| FDS-thenar | 3.5±1.9 – 4.5±2.0 | 3.5±1.9 – 5.3±2.1 | 3.8±1.8 – 5.5±2.8 | 3.9±1.4 – 5.3±3.2 |
| FDI-FDS | 9.6±3.0 – 3.5±1.9 | 8.9±3.8 – 3.5±1.9 | 9.8±5.7 – 3.8±1.8 | 10.3±3.5 – 3.9±1.4 |

### 3.3. Within-muscle coherence

As mentioned in the Methods, within-muscle coherence could only be calculated for the FDI muscle as too few participants exhibited consistent firing activity of enough motor units for the two other muscles. At least one significant coherence value (z-score>1.65) over the delta and alpha band was observed in all the participants, tasks, and sides (Fig. 3).

**Fig. 3.**
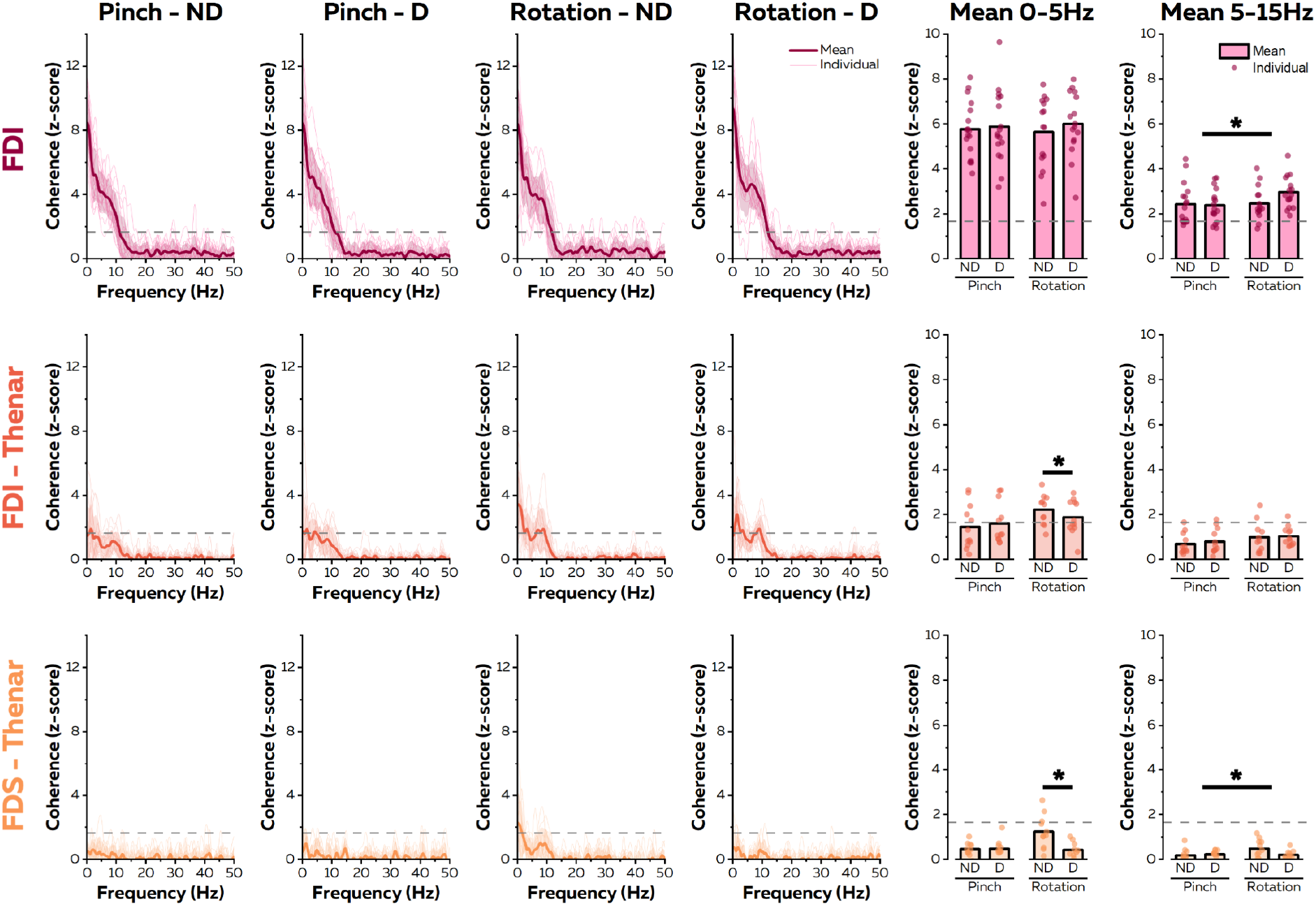
Within and between-muscle coherence. The pooled coherence function is shown for the FDI (within-muscle coherence) and the FDI-thenar and the FDS-thenar muscle pairs (between-muscle coherence). The pooled coherence was calculated from all the combinations (a maximum of 100 random permutations) of two cumulative spike trains, each composed of two motor units. Each thin line corresponds to a participant and the bold line corresponds to the mean across participants. The dashed horizontal dashed line indicates the significant threshold, which was set at 1.65 (95% confidence limit). Individual values of the mean z-score coherence between 0-5 Hz (delta band) and 5-15 Hz (alpha band) are also depicted in the right panels. When considering the delta band, there was a significant interaction between dominance and task. Specifically, a higher coherence was found for the non-dominant than for the dominant hand (regardless of the muscle pair), but this was only observed in the rotation task. When considering the alpha band, there was a main effect of task for both the FDI and the FDS-thenar pair. Only the main statistical results are shown. ND, non-dominant; D, dominant; first dorsal interosseous; FDS, flexor digitorum superficialis.

When considering the mean value of the z-score in the bandwidth 0-5 Hz (*delta* band), we found no significant main effect of task (F(1, 45)=0.001; p=0.981), no significant main effect of dominance (F(1, 45)=0.661; p=0.42), and no significant interaction between task and dominance (F(1, 45)=0.165; p=0.687). When considering the mean value of the z-score in the bandwidth 5-15 Hz (*alpha* band), we found a significant main effect of task (F(1, 45)=7.087; p=0.011), but neither a main effect of dominance (F(1, 45)=2.181; p=0.147) nor an interaction between task and dominance (F(1, 45)=3.395; p=0.072). Specifically, the mean coherence in the *alpha* band was higher for the rotation task (z-score: 2.72±0.75) than for the pinch task (z-score: 2.40±0.81).

### 3.4. Between-muscle coherence

As indicated in the Methods section, coherence could not be calculated for the FDS-FDI pair due to the very low number of motor units that satisfied the analysis criteria. When considering the two intrinsic hand muscles (FDI-thenar pair), at least one significant coherence value (z-score>1.65) was found in the majority of the participants in both the *delta* band (67%, 92%, 100%, and 92% of the participants for pinch/non-dominant, pinch/dominant, rotation/non-dominant, and rotation/dominant, respectively) and the *alpha* band (58%, 67%, 73%, and 92% of the participants for pinch/non-dominant, pinch/dominant, rotation/non-dominant, rotation/dominant, respectively). When considering the FDS-thenar pair, at least one significant coherence value was observed in the *delta* band in the majority of the participants for the rotation task performed with the non-dominant hand (73%), but not for the other tasks (18%, 40%, and 40% of the participants or pinch/non-dominant, pinch/dominant, and rotation/dominant, respectively). A significant coherence value in the *alpha* band was observed in a small proportion of participants (18%, 30%, 46%, and 20% of the participants for pinch/non-dominant, pinch/dominant, rotation/non-dominant, and rotation/dominant, respectively).

Statistical analyses were performed for the FDI-thenar and FDS-thenar pairs (Fig. 3). When considering the mean value of the z-score in the *delta* band, we found a significant main effect of the muscle pair (F(1, 76)=58.525; p<0.001), a significant main effect of the task (F(1, 71)=10.754; p=0.002), and a significant interaction between dominance and task (F(1, 69)=6.009; p=0.017). Specifically, the mean coherence was higher for the FDI-thenar (1.79±0.87) than for the FDS-thenar (0.67±0.59) pair, regardless of the side and task. Also, the mean coherence in the *delta* band was higher for the rotation task (1.74±0.85) than for the pinch task (0.98±0.88; p<0.001) when performed with the non-dominant hand. Furthermore, higher coherence was found for the non-dominant than for the dominant hand, but this was only observed in the rotation task (1.74±0.85 vs 1.23±0.96; p=0.018). No other significant interaction was found (all p>0.285).

When considering the mean z-score in the *alpha* band (Fig. 3), we found a significant main effect of muscle pair (F(1, 74)=52.123; p<0.001), with greater coherence for the FDI-thenar (0.88±0.54) than for the FDS-thenar pair (0.29±0.32). We also found a significant main effect of task (F(1, 70)=8.836; p=0.004), with a higher coherence for the rotation task (0.70±0.57) than for the pinch task (0.50±0.48). There was no significant interaction (all p>0.181). Of note, the same differences were observed when considering only the right-handed participants (data not shown).

### 3.5. Pairwise cross-correlation

To estimate the level of common input between individual motor neurons, we calculated the correlation between their smoothed discharge rates. As this analysis required a minimum of only one motor unit per muscle, it could be performed on a larger number of participants (n=16, 13, 11, 13, 12, and 12 for FDI, thenar, FDS, FDI-thenar, FDS-thenar and FDS-FDI, respectively), and importantly, it could be performed for each muscle/muscle pair. To account for the fact that the strength of common synaptic inputs between two motor neurons is not necessarily proportional to the degree of correlation between their outputs (De la Rocha et al., 2007), we used a conservative approach where only significant correlation coefficients were considered (see Methods). Specifically, we calculated the ratio between the number of significant correlations and the total number of pairwise correlations (Fig. 4).

**Fig. 4.**
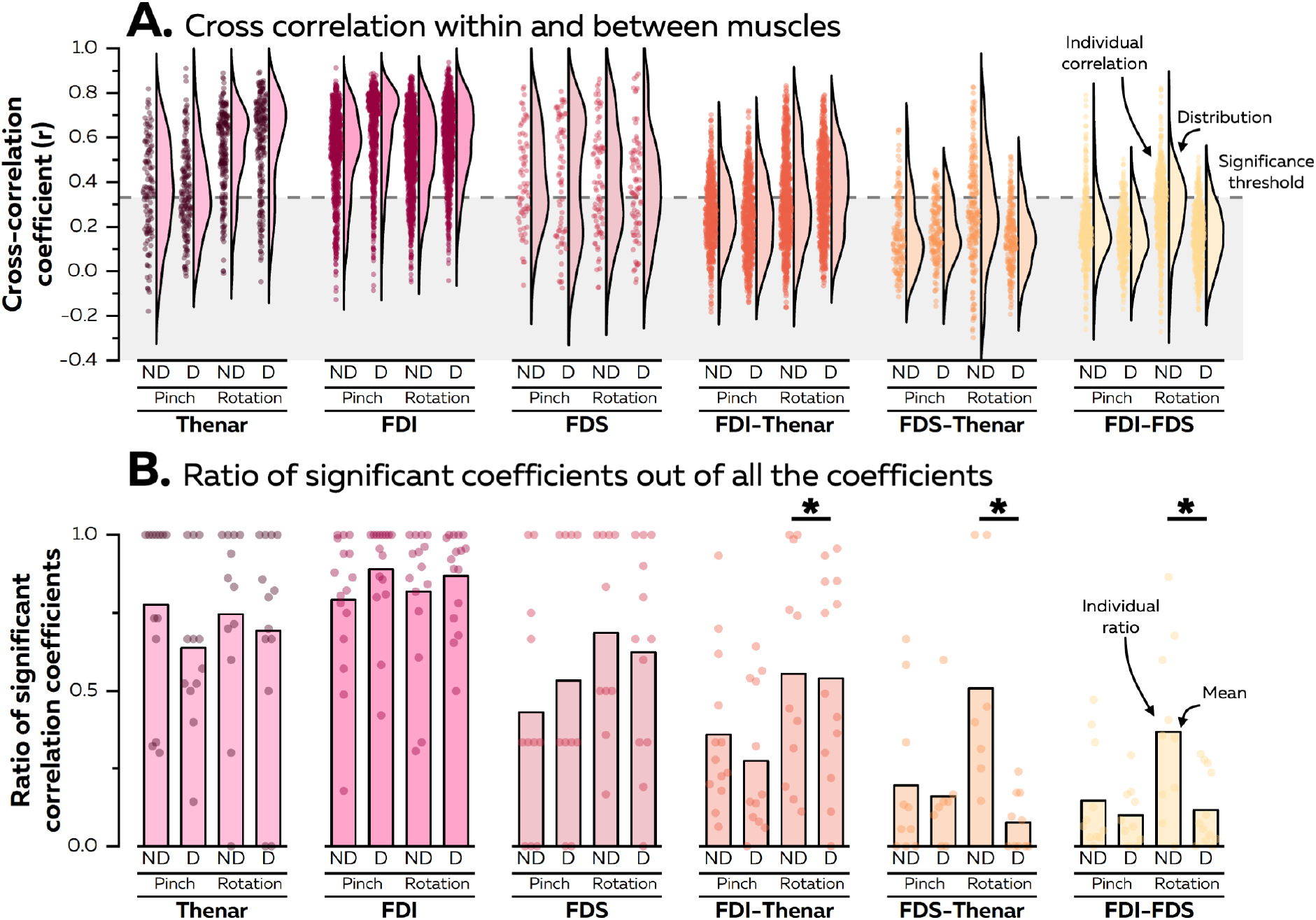
Cross-correlation between smoothed discharge rates. Cross-correlation was calculated for each pair of motor neurons (Panel A). We used a conservative approach where only significant correlation coefficients were considered (Panel B). Specifically, we calculated the ratio between the number of significant correlations and the total number of pairwise correlations. When considering muscle pairs, we found an interaction between task and dominance. Specifically, regardless of the muscle pair, a higher ratio of significant correlations was found for the non-dominant than the dominant hand, but this was observed only in the rotation task. Only the main statistical results are shown. ND, non-dominant; D, dominant; first dorsal interosseous; FDS, flexor digitorum superficialis.

When considering individual muscles (i.e., the correlation between motor neurons from the same pool), we found a main effect of muscle (F(2, 137)=14.537; p<0.001), but no other main effect or interaction (all p>0.121). Specifically, the ratio of significant correlations was higher for the FDI than for both the FDS (p<0.001) and the thenar (p=0.020) (Fig. 4); and higher for the thenar than the FDS (p=0.027).

When considering muscle pairs (i.e., correlation between motor neurons from different pools), we found main effects for muscle (F(2, 109)=12.895; p<0.001), task (F(1, 104)=14.470; p<0.001), and dominance (F(1, 103)=12.394; p<0.001). The ratio of significant correlations was higher for the FDI-thenar than for both the FDS-thenar (p=0.001) and the FDI-FDS (p<0.001) pairs. There was no significant difference between the FDS-thenar and the FDI-FDS (p=0.653) pairs. Furthermore, there was an interaction between task and dominance (p=0.043). Specifically, regardless of the muscle pair, a higher ratio of significant correlations was found for the non-dominant hand than for the dominant hand (p<0.001), but this was observed only in the rotation task. Furthermore, the ratio was higher during rotation than during the pinch task (p=0.001), when these tasks were performed with the non-dominant hand. No other significant interaction was found (all p>0.064).

## 4. Discussion

In this study, we estimated common synaptic inputs across spinal motor neurons from hand muscles during two motor tasks that represent fundamental motor behaviours. We verified that handedness was associated with differences in motor function, as assessed by the Purdue Pegboard Test and the Finger Tapping Tests. Consistent with our hypothesis, our two complementary analyses led to the same conclusion of lower common synaptic input between motor neurons innervating different muscles in the dominant than the non-dominant hand, which was only observed during the isometric rotation task. This lower level of common input might confer greater modularity/flexibility in the recruitment of motor units and, therefore, in the mechanical outputs. Whether this difference between the dominant and non-dominant arm is the cause or the consequence of handedness remains to be determined.

### Common synaptic input between motor neurons that innervate the same pool

Previous studies evaluated common input within muscles of the dominant and non-dominant arm through the correlation of motor unit discharge times over the full bandwidth (referred to as short-term synchronization) (Farina & Negro, 2015). Conflicting results have been reported, with either a higher [extensor carpi radialis (Schmied *et al*., 1994)] or a lower [FDI (Semmler & Nordstrom, 1995)] synchrony in muscles from the dominant than the non-dominant side. Because the input-output nonlinearity of individual motor neurons is more pronounced at higher frequencies (Farina *et al*., 2016), the level of synchronization between trains of action potentials of two motoneurons, which considers the full bandwidth, does not accurately reflect the degree of common synaptic input between these motor neurons (de la Rocha *et al*., 2007). Consequently, the absence of synchronization cannot be taken as conclusive evidence for the absence of common input (Farina & Negro, 2015).

In our study, we used two complementary approaches. First, we performed a coherence analysis on groups of motor neurons, which allowed us to identify common input at different bandwidths. The *delta* band is classically associated with the common drive for control of muscle force (De Luca & Erim, 1994; Negro *et al*., 2009), and the *alpha* band is classically associated with the contribution of the muscle afferents and other spinal circuitries (Williams and Baker, 2009). Notably, this analysis could only be performed for the FDI muscle. We observed a significant level of coherence over the delta and alpha band (z-score>1.65) in all participants, and for all tasks and on both sides. This is consistent with previous work showing a high level of synchrony between motor neurons from the FDI muscle (Datta & Stephens, 1990; Semmler & Nordstrom, 1995). Furthermore, we found no difference between the dominant and non-dominant side, which indicates that handedness is not associated with a different level of common synaptic input when considering motor neurons from the same pool, at least for the FDI muscle. However, when compared to the pinch task, we observed a higher coherence in the alpha band during the rotation task (Fig. 3), which could reflect the need for greater somatosensory and tactile afferent inputs during this task. It should be noted that larger coherence in the alpha band has also been associated with larger force tremor (Laine *et al*., 2014), which is consistent with the larger coefficient of variation of the torque observed during the rotation task when compared to the pinch task. This is an important finding that indicates the sensitivity of our analysis to reveal different neural control strategies between tasks with different mechanical constraints. Therefore, it makes us confident that the lack of difference between sides is not explained by the lack of sensitivity of our approach. We also calculated the correlation between the smoothed discharge rate of individual motor neurons and conservatively considered only the significant correlations. As this analysis required a minimum of two motor units per muscle, it could be extended to the FDS and to the thenar muscles. Overall, this analysis confirmed the absence of an effect of handedness on the level of common input shared between motor neurons from the same pool (Fig. 4).

### Common synaptic input between motor neurons that innervate different pools

Short-term synchronization between motor neurons is often observed between anatomically defined synergist muscles, or between muscles with similar actions, although the level of such synchrony is typically lower than that found within the muscle (Bremner *et al*., 1991; Keen & Fuglevand, 2004). In this way, we observed a significant level of common input between the two intrinsic muscles (FDI and thenar) in the vast majority of the contractions, regardless of whether the analysis (population or individual motor neuron level). This is consistent with previous observations made during tasks that required both muscles to produce force in different directions (Del Vecchio *et al*., 2019). Interestingly, the FDI and thenar can be theoretically activated independently, but they shared a significant level of common input during the two motor tasks that we investigated. Sending common inputs to motor neurons innervating different muscles might be an effective way for the central nervous system to reduce the dimensionality of the control, as suggested by the synergy theory (d’Avella & Bizzi, 2005), and to synchronise their force outputs (Santello & Fuglevand, 2004). In addition, common input control implies the orderly recruitment of motor neurons, as the group of motor neurons receiving common input can only be recruited according to the Henneman’s size principle (De Luca & Erim, 1994). Taken together, this suggests that, despite achieving the important goal of reducing the control dimensionality, the presence of common inputs may reduce the ability to make fine motor adjustments. This is particularly important to consider to properly interpret differences between sides, as discussed below.

It is noteworthy that regardless of the analysis, a less common input was observed between intrinsic (thenar or FDI) and extrinsic (FDS) muscles (Fig. 3 and 4), which is consistent with previous observations made on the same muscles and tasks using interferential surface EMG signals (Laine & Valero-Cuevas, 2017). It is also consistent with previous results suggesting that motor neurons from distant muscles may receive common inputs, but in a smaller proportion than that observed between neighbouring synergist muscles (Gibbs *et al*., 1995).

To the best of our knowledge, the effect of hand dominance on the level of common synaptic input between motor neurons innervating different pools has received limited attention. Hockensmith et al. (2005) found that the magnitude of synchrony between two extrinsic hand muscles (flexor pollicis longus and flexor digitorum profundus) was markedly higher in the dominant than in the non-dominant hand. These findings contrast with our results showing either no difference between sides when considering the isometric pinch task, or a lower level of common input for the dominant side when considering the rotation task, regardless of the muscle pair. The discrepancy between our results and those from Hockensmith et al. (2005) might be explained by the fact that they focused on different muscles (two extrinsic hand muscles vs. intrinsic and extrinsic muscles in our study) or by the characteristics of the task (simple pinch task performed at a very low intensity). Of note, the lower level of common input observed in our study during the rotation task was not associated to a difference in force variability between sides. As mentioned above, this lower level of common input may allow more flexibility in the recruitment of motor units and, therefore, in mechanical outputs. In this way, Rossato et al. (2022) observed a redistribution of neural drive across synergist muscles during a fatiguing task, but this was only observed between muscles that shared a low level of common drive. In addition, our results are well in line with previous work, which investigated handedness in precision tasks from force signals, and which showed that the thumb and index finger of the dominant hand might be more independently controlled by the central nervous system than the non-dominant digits (Reilly & Hammond, 2004; Li *et al*., 2015).

The fact that a dominance effect was only found for the isometric rotation task may be explained by specific mechanical constraints, which required our participants to both pinch and rotate a dial. Based on the previous observation of differential changes in the behaviour of FDI motor units with changes in force direction (Desmedt & Godaux, 1981), overcoming these specific task constraints may have been facilitated by a more independent control of motor neurons. Because rotating a dial is almost always performed with the dominant hand in daily life, an optimal level of common inputs may have been fine-tuned with task repetition. This is consistent with previous work reporting a fractionation of synergies (i.e. an increase in the number of common inputs) during development (Dominici *et al*., 2011; Cheung *et al*., 2020). However, we cannot exclude an alternative explanation whereby the lower common inputs observed in the dominant hand are the cause rather than the consequence of handedness.

### Methodological considerations

This study requires the consideration of three main methodological aspects. First, despite the fact that the borders of the FDS muscle were located with B-mode ultrasound prior to the electrode placement, we cannot ascertain the selectivity of the measurements to the FDS motor units, as the borders of the grid were likely close to neighbouring muscles. It is therefore possible that the electrode picked up activity from neighbouring muscles. However, it is likely that if it happened, it did not differ between sides, which makes us confident in our main conclusion about the effect of dominance on the common input involving the FDS muscle.

Furthermore, even though cross talk can greatly affect interferential EMG signals (Germer *et al*., 2021), it is not a major issue when identifying motor units. Indeed, only a small proportion (< 1%) of motor units were identified as ‘cross-talk’ units in previous studies that verified that the identified motor units did originate from the target muscles on which the grid was placed and did not originate from crosstalk from a neighbouring muscle (Hug *et al*., 2021c; Rossato *et al*., 2022). Second, despite the fact that we identified a similar (or even a higher) number of motor units than in previous HDsEMG studies that targeted the same muscles (Del Vecchio *et al*., 2019; Tanzarella *et al*., 2021), we failed to perform each analysis for all of our participants. However, we used two complementary approaches, which, taken together, allowed us to strengthen our main conclusions. Finally, we tested only three left-handed participants. Although this represents the proportion of left-handlers in the general population, it is important to note that certain features of the pyramidal tract projecting to the right side of the spinal cord tend to be different from those projecting to the left, regardless of handedness (Nathan *et al*., 1990). As such, some of the differences that we observed might be related to side preference rather than hand preference.

## Conclusions

By identifying the behaviour of single motor neurons over three hand muscles, we demonstrated that there is a reduced common synaptic input between motor neurons innervating different muscles in the dominant compared to the non-dominant hand. This could confer greater flexibility in the recruitment of motor units to comply with task constraints. Further research is needed to determine whether the difference between the dominant and non-dominant arm is the cause or the consequence of handedness.

## Acknowledgements

François Hug and Antoine Nordez are supported by a fellowship from the *Institut Universitaire de France* (IUF). Support was received from the IFTH (*Institut Français Textile et Habillement*). The authors thank Aurélie Sarcher for her help during the experiments.

